# Poly(ADP-ribose)polymerase1 facilitates the nucleosome disassembly

**DOI:** 10.64898/2026.01.21.700758

**Authors:** A.A. Alekseev, M.M. Kutuzov, E.A. Belousova, I.D. Goncharov, A.A. Vasileva, M.A. Khodorkovskii, O.I. Lavrik

## Abstract

Being the basic building blocks of chromatin, nucleosomes and their stability determine the genome accessibility for different DNA-dependent proteins. This characteristic is labile under all cell-life processes. One of the abundant DNA-binding proteins, which is important for genome compaction, is poly(ADP-ribose)polymerase1 (PARP1). Despite the extensive experimental data on the chromatin compaction regulation under ADP-ribosylation, the details of the interplay of nucleosome with PARP1 in the absence of protein activation remain unclear. In this study, we analyzed the changes in the nucleosome wrapping upon PARP1 interaction using a single-molecule approach — optical tweezers. We demonstrate that PARP1 binding leads to weakening of the contacts that support the nucleosome core.

## Introduction

The chromatin organization is required for DNA compaction as well as the regulation of the DNA-dependent processes [Mansisidor and Risca, 2022]. The basic unit of chromatin is the nucleosome core particle (NCP), whose compaction is under fine regulation. There are two main mechanisms providing correct DNA compaction: enzymatic, which includes histone posttranslational modifications and the action of ATP-dependent nucleosome remodeling, and the second one is the action of non-histone chromatin proteins changing NCP conformations via binding to them [Zhou et al., 2019].

PARP1 is best known as a central regulator of DNA damage detection and repair [Chaudhuri and Nussenzweig, 2017], can affect the NCP structure. It is a clinically validated target, with several approved molecules that are particularly effective in the treatment of cancers bearing different mutations leading to the falling down of RAD51 functionality [Jackson and Moldovan, 2022; O’Malley et al., 2023; Peng et al., 2025]. Multiple studies describe the main PARP1-mediated mechanism of regulation of NCP compaction via histone poly(ADP-ribosyl)ation. At the same time, the binding of PARP1 to the NCP affects nucleosome compaction without activation [Sultanov et al., 2017]. There are some data demonstrating the ability of PARP1 to interact with NCP like a linker histone H1 [Krishnakumar et al.,2008]. It has been shown that histone H1 does not influence NCP compaction density but reduces its dynamics toward the stabilization of a defined conformation [Bednar et al., 2017; Würtz et al., 2019]. In terms of the set of non-covalent bonds dynamically organizing the NCP structure, the existence of several spatial conformations of NCP can be predicted. Moreover, each of them could be characterized by different relation strength [Armeev et al., 2025; Chen et al., 2024; Hayes and Hansen, 2002]. In this work, using optical tweezers we estimated the strength of NCP contacts under its binding by PARP1.

## Results and discussion

Optical tweezers are a common method for measuring the NCP strength, first used to measure the unwrapping force of individual NCP [Bennink et al., 2001]. It should be noted that the average meaning of NCP unwrapping force can vary in a wide range depending on the DNA model used [Widom, 2001]. This is mainly based on electrostatic differences of different nucleosome positioning DNA sequences (NPS) and histone variant sets, as well as the influence of buffer components.

In our work we used a model system based on Widom’s clone 601 with eight tandem NPSs. First, we defined the typical force-extension profile of the specified DNA probe and determined the force for individual NCP unwrapping events (Fig. 1A).

**Figure 1.**
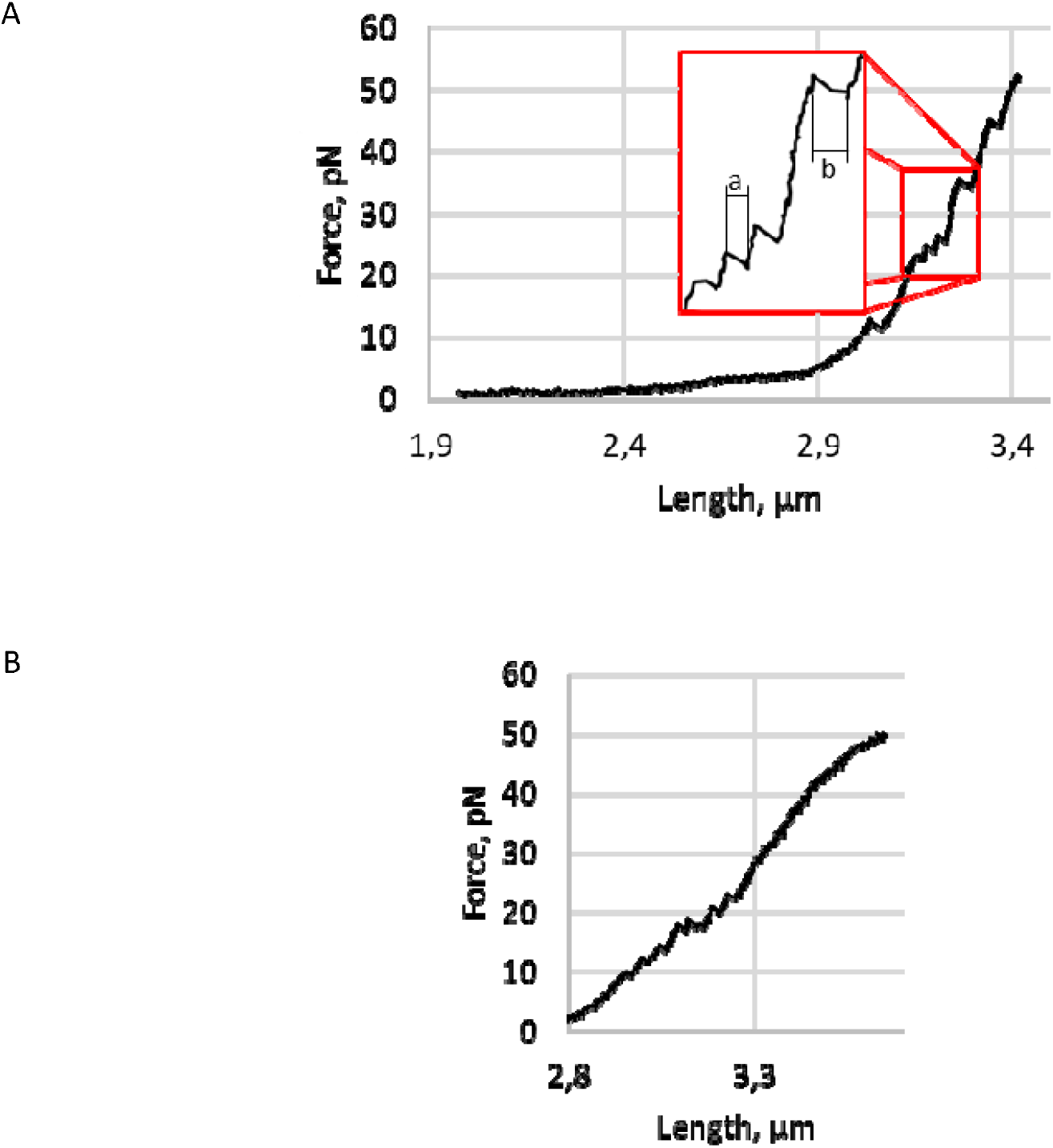
The force-extension profile of NCP unwrapping. Typical force profile demonstrating the dependence of tensile force on NCP-containing DNA probe stretching without PARP1 (A) and with 50 nM PARP1 (B). The characteristic stretching lengths of disassembly of a single (a) and simultaneously two NCPs (b) are 20 and 40 nm, respectively.

The observed force-extension profile of the NCP turned out to correspond to the literature data and exhibited two principal regions [Díaz-Celis et al., 2022]. One of those has a slight gradual rise of characteristic rupture force in diapason at approximately 2.55-2.70 μm. It corresponds to the release of the outer DNA turn of the NCP. Another one has a few “teeth” on the curve and is compliant with the unwinding of the inner DNA turn from the NCP core.

In Fig. 1A(a), the characteristic “teeth” corresponding to the unwrapping of the NCP inner turn led to an elongation of the DNA probe by 20 nm, which corresponds to ∼70 base pairs, and are clearly visible in the force range of approximately 10-45 pN. Several “teeth” value unwrapping is about 40 nm, which most likely corresponds to unwrapping of two NCPs simultaneously with the same force value (Fig 1A(b)).

The protocol of preparing an NCP-containing DNA probe does not exclude the possibility of forming complexes on non-specific sequences, allowing different numbers of nucleosomes into one DNA molecule. Analyzing the force-extension curve of the DNA probe, we defined the number of NCP per DNA molecule. The distribution had a maximum at seven-eight (Fig. S1). Next, we combined the values of the forces obtained in the DNA stretching experiments. The distribution is presented on histograms in Fig. 2.

**Figure 2.**
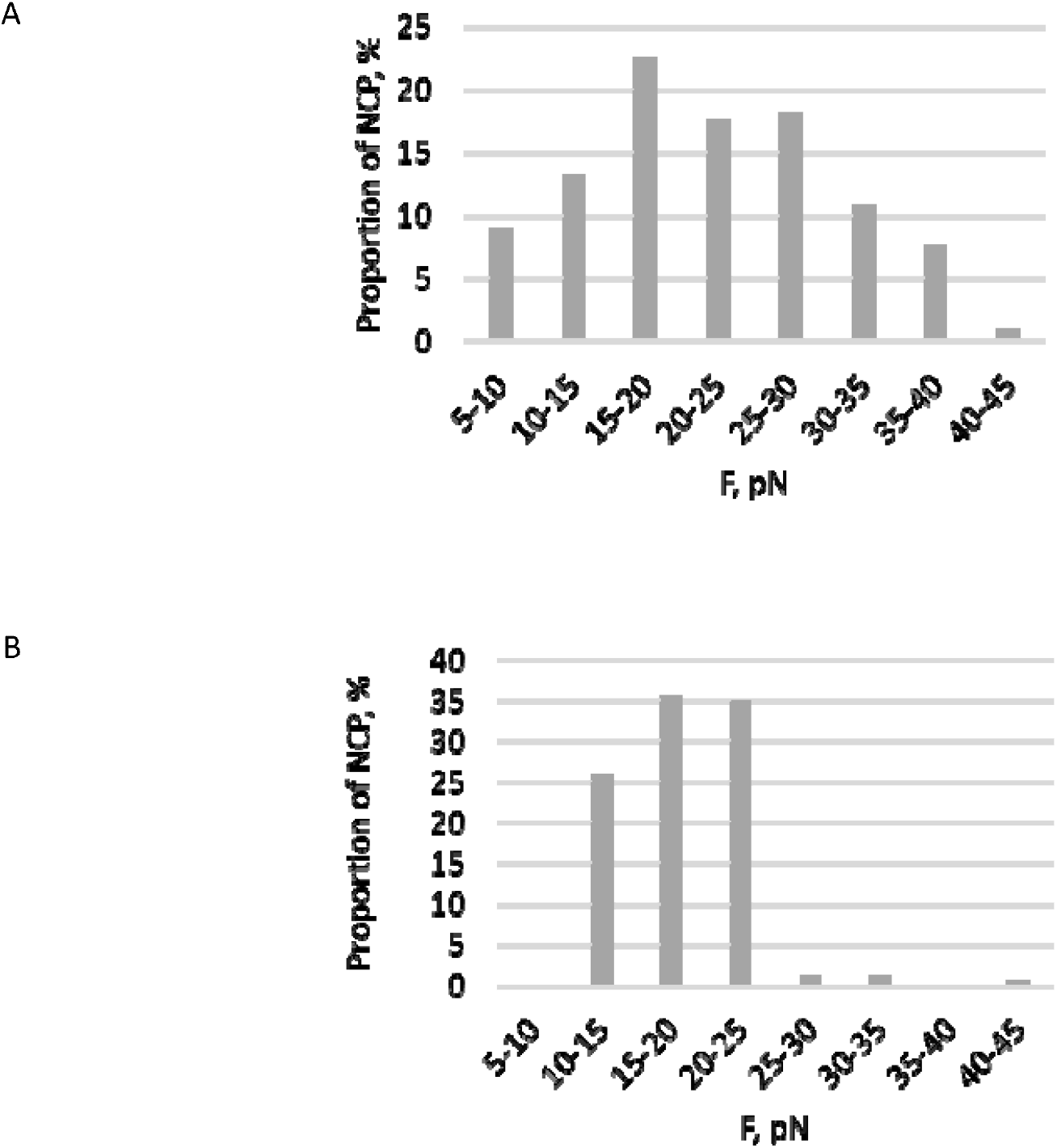
The distribution of NCP unwrapping force. The histograms display the distribution of the amount of NCP complexes by strength tension of probes without PARP1 (A) and with PARP1 (B). The force ranges of disassembly are on the X-axis, in pN. The representation of the NCP complexes at a defined strength range is on the Y-axis, in percentage. The number of NCP in the calculation is three hundred fifty-eight (A) and one hundred fifty-six (B).

The data in Fig. 2A correlates to the literature [Mihardja et al., 2006]. The range of forces required for NCP unwrapping varies in the range of 5 - 45 pN. The majority of events of NCP unwrapping occurred at tensile forces in the range of 15–30 pN.

To define the impact of PARP1 on the NCP tension profile, we performed similar experiments using a freshly assembled DNA in the presence of 50 nM PARP1 (Fig. 1B). The typical tension profile of NCP disassembly in this case is generally similar to the DNA probe tension in the absence of PARP1 (Fig. 1A) and characterized by a few “teeth” of NCP unwrapping; however, the signature of the “teeth” is different. In particular, these distinctions are expressed in the lower force magnitude corresponding to each “tooth.”

The distribution of the NCP unwrapping force in the presence of PARP1 is shown in Fig. 2B. It is evident that the distribution differs from that shown in Fig. 1B by the virtual absence of nucleosomes, for which the unwrapping force is 25-45 pN and whose contribution to the distribution in the absence of PARP1 was more than 35%.

PARP1 has previously been shown to bind [Sukhanova et al., 2016] and condense naked DNA [Bell et al., 2021] under sub-piconewton forces diapason due to the stabilization of DNA loops. We observed also a noticeable difference of DNA force extension at tensile forces greater than 15 pN. The curve with PARP1 is less steep, indicating a decrease in DNA stiffness in the presence of this protein (Fig. 3). This phenomenon likely indicates the existence of an alternative DNA state during PARP1 interaction. It could be an additional argument for the multifaceted interaction of PARP1 with the nucleosome.

**Figure 3.**
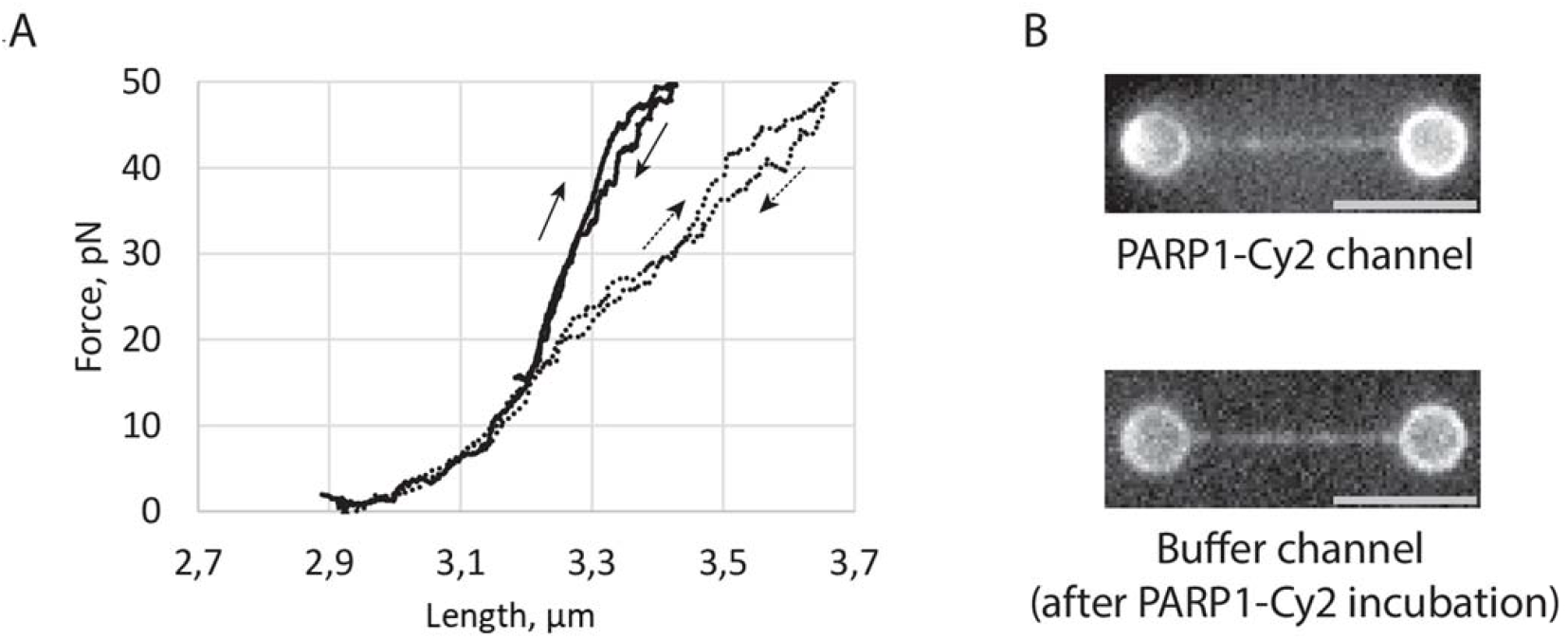
Interaction of PARP1 with DNA. (A) Force-extension curves of a 9.9 kbp DNA fragment without NCP in buffer (solid line) and in the presence of 50 nM PARP1 (dotted line). (B) Single-molecule fluorescence imaging of PARP1-Cy2 binding to a representative DNA molecule from the 22-24 kbp preparation (see Materials and methods for details). Top: DNA in the microfluidic channel containing 50 nM PARP-Cy2; bottom: the same DNA molecule after transfer to a buffer channel lacking free PARP1-Cy2. Scale bar, 5 μm.

There are abundant data about the influence of PARP1 on the NCP state in the context of genome compaction. At the beginning, intriguing results allowed us to hypothesize the competition between PARP1 and linker histone H1 for NCP binding [Kaiser et al., 2020; Koshkina et al., 2025; Shan et al., 2014; Sharma et al., 2019]. The recent studies even proposed the complex model of interaction of PARP1 with NCP in the linker region [Clark et al., 2012; Koshkina et al., 2025]. The smFRET-based studies disclosed distance changes between neighboring gyres of nucleosomal DNA in the axial direction under PARP1 binding that could lead to distortion of the canonical NCP conformation [Sultanov et al., 2017]. In spite of that, our AFM study of the NCP compaction displays no significant changes of DNA arms positioning in the radial direction upon PARP1 binding [Ukraintsev et al., 2023]. These contractionary results could be a consequence of the approach’s limitations: the AFM technique suggests the molecular adsorption on the mica surface and could be a failure to detect the axial distortions of DNA into NCP. Thus, the results obtained by the smFRET and AFM assays could be considered as supplementing each other. Additionally, PARP1 has recently been shown to demonstrate higher affinity to partially unwrapped NCP over the canonical one [Schaich et al., 2026]. Moreover, the interplay between the functioning of PARP1 and chromatin remodeling factors was shown. For example, ATP-dependent factor ALC-1 exhibits the cooperative action to PARP1 on NCP under PARP1 activation [Gottschalk et al., 2012]. Further, the details remain to be determined, but there are some evidences that PARP1 and its interaction with the nucleosome core particle play a role in ALC1-dependent nucleosome remodeling [Ooi et al., 2021]. Thus, together with our results, it may suggest a potential role of PARP1 in the reorganization of NCP in a deeper stage when the NCP is not already fully wrapped.

## Conclusions

In this study, using optical tweezers, we demonstrated the changes in the nucleosome core wrapping tension in the presence of PARP1. Overall, the available data suggest that the PARP1 interaction with the nucleosome is complicated. Outside the context of DNA damage, PARP1 may act as a regulator of nucleosome core density both directly and by interaction of PARP1 with linker DNA regions, like histone H1, leading to stabilization of the core particle. Alternatively, PARP1 may interact directly with nucleosomal DNA and reduces the energy barrier at the loosening or unwrapping of the nucleosome core. This potential mechanism agrees well with the monkey bar model proposed earlier for the interaction of PARP1 with DNA [Rudolph et al., 2018].

Moreover, it is possible that the site of PARP1 binding affects not only the density of the nucleosome core itself but also the influence on the specific proteins recruitment that mediate nucleosome remodeling or sliding. Since nucleosomes undergo remodeling under DDR to implement an alternative gene expression, PARP1 could be one of the regulators of stress-susceptible or stress-resilient factors in the implementation of DNA-dependent processes.

## Matherials and methods

### DNA constructs and proteins

Human PARP1 and chicken histones are purified as described earlier [Amé et al., 2011; Kutuzov et al., 2019]. The 9.9 kbp fragment bearing eight tandem Widom’s 601 sequences separated by 25 bp linkers was released by XbaI digestion from the pKYB1-based plasmid containing the corresponding 8×NPS inserts [Spakman et al., 2020]. The linearized plasmid was then ligated with a 50-fold molar excess of biotinylated oligonucleotide (5’-CTAGCGAGTGXXXXX-3’, X denotes biotin tag) in the presence of a short helper oligonucleotide (5⍰-CACTCG-3⍰) to facilitate efficient ligation to the XbaI overhang [Horspool et al., 2010]. Unincorporated oligonucleotides were removed by gel-filtration through a Bio-Spin P-30 column (Bio-Rad). Nucleosomes were then reconstituted onto the resulting biotinylated DNA construct as described in [Kutuzov et al., 2019] with minor variations. Definitely, we do not perform here the pre-reconstitution of NCP and verify the efficiency of NCP preparation by optical tweezers instead of native gel-electrophoresis.

For single-molecule fluorescence experiments, a set of linear DNA molecules of 22 kbp and 24 kbp was prepared from bacteriophage λ DNA (48.5 kbp) as previously described [Alekseev et al., 2022]. The DNA was nucleosome-free, double-stranded, and contained no intentionally introduced damage.

Fluorescent labelling of PARP1 on the terminal amino group using succinimidyl esters of 5(6)-carboxycyanine2 (Cy2-SE) was performed as described [Moor et al., 2015]. The stoichiometry of protein labelling did not exceed 1 mole of dye per mole of protein.

### Optical tweezers setup

Optical trapping experiments were performed using a custom dual-trap instrument built around a 1064 nm Nd:YVO_4_ CW laser (5 W, Spectra-Physics BL-106C) and a high-NA oil-immersion objective (LOMO 100×, NA 1.25), essentially as detailed previously [Alekseev et al., 2020; Alekseev et al., 2022]. One trap was steered with sub-nanometer precision using a piezo-driven mirror (Physik Instrumente S-330.80L). Trap stiffness was determined by the viscous-drag method on a high-precision piezo stage (P-561.3DD). Real-time force and bead-to-bead distance were recorded at 33 Hz via bead-tracking in LabVIEW. Measurements were carried out in a five-channel microfluidic chamber (Lumicks).

### Single-molecule assay

All single-molecule experiments were performed at 22°C in buffer containing 20 mM Hepes-NaOH (pH 7.5), 100 mM NaCl, 2 mM MgCl_2_, 0.2% (w/v) BSA, and 0.02% (v/v) Tween-20. Microfluidic channels were blocked with 0.5% (w/v) Pluronic F-127 and 0.1% (w/v) BSA [Belan et al., 2021]. One of the five channels of the microfluidic chamber was filled with 0.01% (w/v) 2.1 μm streptavidin-coated polystyrene beads (Spherotech), and the neighbouring channel contained 50 pM nucleosome-array DNA in measurement buffer. Two beads were optically trapped in the bead channel, moved into the DNA channel, and a single DNA molecule was tethered between them. Successful tethering and the presence of multiple nucleosomes were confirmed by the characteristic-force rip pattern. Force-extension curves were recorded in position-clamp mode by increasing the distance between the traps at 0.1 μm s^−1^ while monitoring bead positions and force with 30 ms time resolution.

When the impact of PARP1 on NCP was estimated, the DNA tether adjusted to a tension of 2-3 pN, and keeping the trap positions fixed, the tethered DNA was rapidly moved into the adjacent channel containing 50 nM PARP1 in measurement buffer. After 1 min incubation at the tension 2-3 pN, the force-extension curve was recorded in the channel with PARP.

Single-molecule fluorescence imaging of PARP1-Cy2 binding to trapped DNA molecules was performed using essentially the same acquisition settings and optical configuration as previously described [Alekseev et al., 2022]. Briefly, fluorescence was excited with a 473 nm continuous-wave laser (100 mW) attenuated using a 2.0 OD neutral density filter. Images were acquired with an EMCCD camera at 100 ms exposure time and an electron multiplier gain of 3000 and stored as uncompressed TIFF files. During imaging, the DNA was maintained under mild tension of ∼3 pN to keep it extended and prevent potential PARP1-induced DNA condensation. Images were processed in FIJI ImageJ. Background illumination gradients were corrected by local background subtraction. Brightness and contrast were adjusted linearly. Unprocessed (raw) images are provided in Fig. S2 and Fig. S3.

### Calculations

The measured forces of NCP disassembly were sorted into the groups with force intervals of 5 pN. The histograms display the representation of each group as a percentage of the total sample.

## Acknowledgments

We thank Georgii Pobegalov (University College London and The Francis Crick Institute, London, UK) for stimulating discussions on experimental design.

## Funding

The work was performed in financial support of RSF №24-74-00107 (single molecule experiments), RSF №25-74-30006 (protein purification and nucleosome probe preparation), and the Russian state-funded project for the ICBFM SB RAS (grant No. 125012300658-9) (for O.I. Lavrik).

## Notes

### Competing Interest Statement

The authors have declared no competing interest.

