## Supplemental Fig. S1 for "Poly(ADP-ribose)polymerase1 facilitates the nucleosome disassembly"

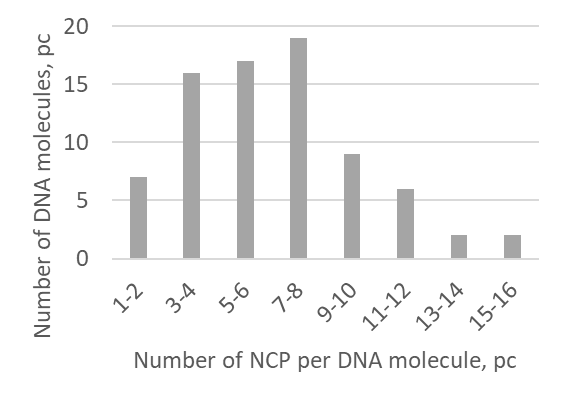

**Figure S1.** The distribution of the NCP number per DNA probe. The number of DNA molecules in the calculation is seventy-eight.
