## Supplementary figures and images for "Poly(ADP-ribose)polymerase1 facilitates the nucleosome disassembly"

### Supplemental Fig. S2

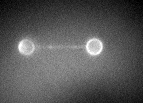

### Supplemental Fig. S3

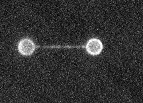
